# A network-based data-mining approach to investigate indole-related microbiota-host co-metabolism

**DOI:** 10.1101/602458

**Authors:** Ana Luisa Neves, Andrea Rodriguez-Martinez, Rafael Ayala, Joram M Posma, MR Abellona U, Julien Chilloux, Jeremy K Nicholson, Marc-Emmanuel Dumas, Lesley Hoyles

## Abstract

**Motivation:** Indoles have been shown to play a significant role in cardiometabolic disorders. While some individual bacterial species are known to produce indole-adducts, to our best knowledge no studies have made use of publicly available genome data to identify prokaryotes, specifically those associated with the human gut microbiota, contributing to the indole metabolic network.

**Results:** Here, we propose a computational strategy, comprising the integration of KEGG and BLAST, to identify prokaryote-specific metabolic reactions relevant for the production of indoles, as well as to predict new members of the human gut microbiota potentially involved in these reactions. By identifying relevant prokaryotic species for further validation studies *in vitro*, this strategy represents a useful approach for those interrogating the metabolism of other gut-derived microbial metabolites relevant to human health.

**Availability:** All R scripts and files (gut microbial dataset, FASTA protein sequences, BLASTP output files) are available from https://github.com/AndreaRMICL/Microbial_networks.

**Contact:** ARM; LH.

## INTRODUCTION

The gut microbiome communicates with its human host through a range of signalling metabolites that bind human targets and impact their signalling pathways, thus modulating human health (Nicholson, Holmes et al. 2005, Holmes, Kinross et al. 2012, Chilloux, Neves et al. 2016). Among these metabolites, indoles – heterocyclic compounds produced upon bacterial degradation of tryptophan or plant metabolism – appear to be particularly relevant for human health. In the last three decades, metabonomic studies have consistently identified indole-containing molecules in human biofluids (Bales, Higham et al. 1984, Danaceau, Anderson et al. 2003, Duranton, Cohen et al. 2012, Bouatra, Aziat et al. 2013). Indoles have been linked to deleterious phenotypes both *in vitro* and *in vivo*. For instance, in endothelial cells, 3-indoxylsulphate induced oxidative stress by modifying the balance between pro- and anti-oxidant mechanisms, stimulating the release of endothelial microparticles and diminishing endothelial-healing ability (Faure, Dou et al. 2006). 3-Indoxylsulphate also aggravates cardiac fibrosis and cardiomyocyte hypertrophy (Yisireyili, Shimizu et al. 2013). In clinical studies, levels of 3-indoxylsulphate seem to be a powerful predictor of overall and cardiovascular mortality (Barreto, Barreto et al. 2009). Another indole, indole-3-propionate, is involved in inflammatory mechanisms and in the maintenance of intestinal barrier integrity (Venkatesh, Mukherjee et al. 2014). The production of indoles depends on gut bacterial action, and therefore is affected by microbiome interventions such as the use of antibiotics (Sun, Schnackenberg et al. 2013) or functional foods (Yang and Tarng 2018). A number of bacterial species involved in different indole synthesis pathways have been described in the literature. Tryptophan can be directly converted to indole by tryptophanase, a bacterial lyase present in *Bacteroides thetaiotaomicron, Proteus vulgaris* and *Escherichia coli* (Jean and DeMoss 1968, Wesoly and Weiler 2012); indole is further conjugated in the liver to form 3-indoxylsulphate. Alternatively, tryptophan can be deaminated by *Clostridium* spp. and *Lactobacillus* spp., producing indole-3-pyruvate, indole-3-lactate, indole-3-acetate and 3-methylindole (Mohammed, Onodera et al. 2003, Attwood, Li et al. 2006, Whitehead, Price et al. 2008, Wikoff, Anfora et al. 2009, Wesoly and Weiler 2012, Russell, Duncan et al. 2013). Studies performed in gnotobiotic mice demonstrated that the production of another indolic compound, indole-3-propionate, was completely dependent on colonisation by *Clostridium sporogenes* (Wikoff, Anfora et al. 2009).

While a number of studies have evaluated the contribution of individual species to the formation of specific indoles, there is a need for a systematic approach to investigate indole synthesis in the context of the gut microbiota ecosystem. In this study, we have developed a computational workflow to predict the contribution of the gut microbiota to the production of disease-related metabolites, here exemplified for indoles. In particular, we used the Kyoto Encyclopaedia of Genes and Genomes (KEGG) (Kanehisa and Goto 2000), a popular reference database comprising biological pathways and cellular processes, as a starting point to identify the metabolic reactions relevant for the production of indoles and the bacterial species involved in these reactions. We then used a BLASTP-based strategy (Altschul, Gish et al. 1990) to predict novel key gut microbial players involved in this metabolic network, offering new insights into the prokaryotic metabolism of tryptophan.

## MATERIALS AND METHODS

### Indole metabolic network

The MetaboSignal R/Bioconductor package (v.3.8) was used to build a KEGG-based tryptophan metabolic network (Rodriguez-Martinez, Ayala et al. 2017). This network has a bipartite structure, where nodes represent two different biological entities: metabolites and reactions. A reaction-to-metabolite distance matrix was generated to identify reactions directly connected to indole-containing compounds. The tryptophan network was then filtered to include only reactions involved in indole production from tryptophan. The reactions from the indole synthesis network were linked to their corresponding organism-specific genes using a two-step procedure: 1) from reaction to KEGG ortholog (KO); 2) from KO to organism-specific genes. The indole synthesis network was further filtered, by excluding reactions which are not undertaken by human or prokaryotic organisms. Three reactions (“rn:R00681”, “rn:R01971”, “rn:R01973”) were included in the network despite not being linked to a KO, because there is evidence that these reactions can occur in some bacterial species (Jean and DeMoss 1968, Hornemann, Hurley et al. 1971, Speedie, Hornemann et al. 1975, Roberts and Rosenfeld 1977, Takai, Ushiro et al. 1977, Du, Alkhalaf et al. 2015). This network was visualized and customized in Cytoscape (v.3.4) (Shannon, Markiel et al. 2003).

### Gut microbial species

A dataset of gut microbial organisms was created by merging the list of gastrointestinal isolates reported at the Human Microbiome Project (HMP) Catalog (Turnbaugh, Ley et al. 2007) with additional species identified in cultivation studies (Rajilic-Stojanovic and de Vos 2014, Browne, Forster et al. 2016, Lagier, Khelaifia et al. 2016). Data from HMP were downloaded on 12 November 2018; and only isolates with complete sequences and annotation were included. The taxize R/CRAN package (v.0.9.6) (Chamberlain, Szoecs et al. 2018) was used to retrieve the taxon identifier (ID) as well as the phylum of each organism.

### BLAST analysis

For each KO involved in indole metabolism, a FASTA file of concatenated prokaryotic protein sequences was generated. These FASTA files were used to run BLASTP searches against the non-redundant (rn) protein sequence database on 30 November 2018. BLASTP results were filtered using an E value cut-off of 1×10^−3^ and 90 % sequence coverage; only hits with ≥ 95 % identity were retained.

### Visualization of results

The results of the analysis were visualized using different approaches. For 3-indoxylsulphate, relevant bacterial species were grouped by phylum and visualized as a network in Cytoscape. For indole-3-acetate, a reaction-species heatmap color-coded according to the ability of a given bacterium to perform each reaction was generated. In order to highlight the most relevant species, the reaction-species matrix was filtered as follows: 1) three different paths leading to the production of indole-3-acetate from tryptophan were identified; 2) each reaction was assigned a value (reaction-value) reflecting its contribution to the corresponding path (i.e. 1/total number of reactions in the path); 3) each species was given a score (Species Contributor Score, SCS), calculated as the sum of the individual reaction-values.

## RESULTS

### Contribution of human and prokaryotic metabolic reactions to the production of indolic compounds

We generated a KEGG-based network illustrating the metabolic reactions involved in the production of indoles from tryptophan. The network was filtered to include only those metabolic reactions that can be performed by human or prokaryotic organisms. Based on this, 16 metabolites and 19 metabolic reactions (8 prokaryotic-specific, 5 human-specific, and 6 present in both human and prokaryotes) were retained (**Figure 1**). In total, 39 % of the reactions of the network are performed exclusively by bacterial and archaeal species, reflecting the relevance of prokaryotes on the metabolism of indoles. Indole-3-pyruvate is the only indolic compound produced upon human metabolism exclusively, by action of the L-amino-acid oxidase. The production of all remaining indoles requires prokaryotic metabolism, either for all reactions of a given path (indole and indole-3-acetate via indole-3-acetamide) or at least for one of them (indoxyl and indole-3-acetaldehyde). Bacterial metabolism is also required for the synthesis of indole-3-lactate, 3-indoleglycolaldehyde and 3-methylindolepyruvate, although these reactions have not been fully characterised yet (i.e. the reactions are not linked to a KO ortholog).

**Figure 1.**
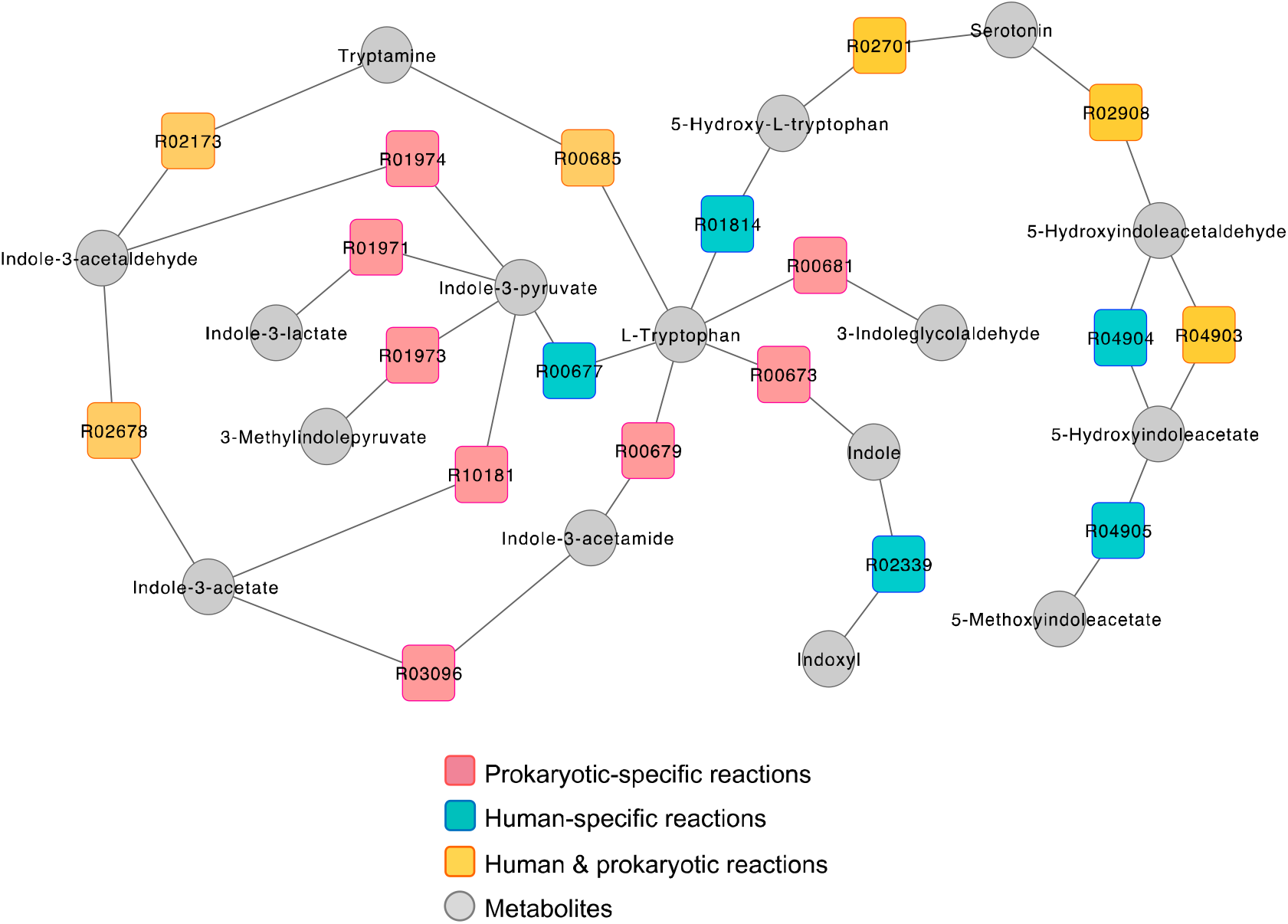
Overview of the production of indoles from tryptophan. Network illustrating the metabolic reactions involved in the synthesis of indoles from tryptophan, where all reactions are irreversible. Only reactions undertaken by human or microbial species are shown. Network nodes represent two different biological entities: reactions (squares) and metabolites (circles).

### BLASTP-based strategy to predict novel key prokaryotic genes and species involved in mammalian-microbial co-metabolism

Next, we used a BLASTP-based strategy to augment the identification of bacterial species involved in indole production reported in KEGG. This approach allowed the identification of 2,626 novel prokaryotic species potentially implicated in indole metabolism (+63.4 % compared to KEGG) (**Figure 2a**). For individual reactions, the increases in the number of prokaryotic species ranged from 7 (“K11816”) to 1,380 (“K00128”) (**Figure 2b**). Similarly, we confirmed that human-specific reactions reported in KEGG were not performed by prokaryotes according to BLASTP searches.

**Figure 2.**
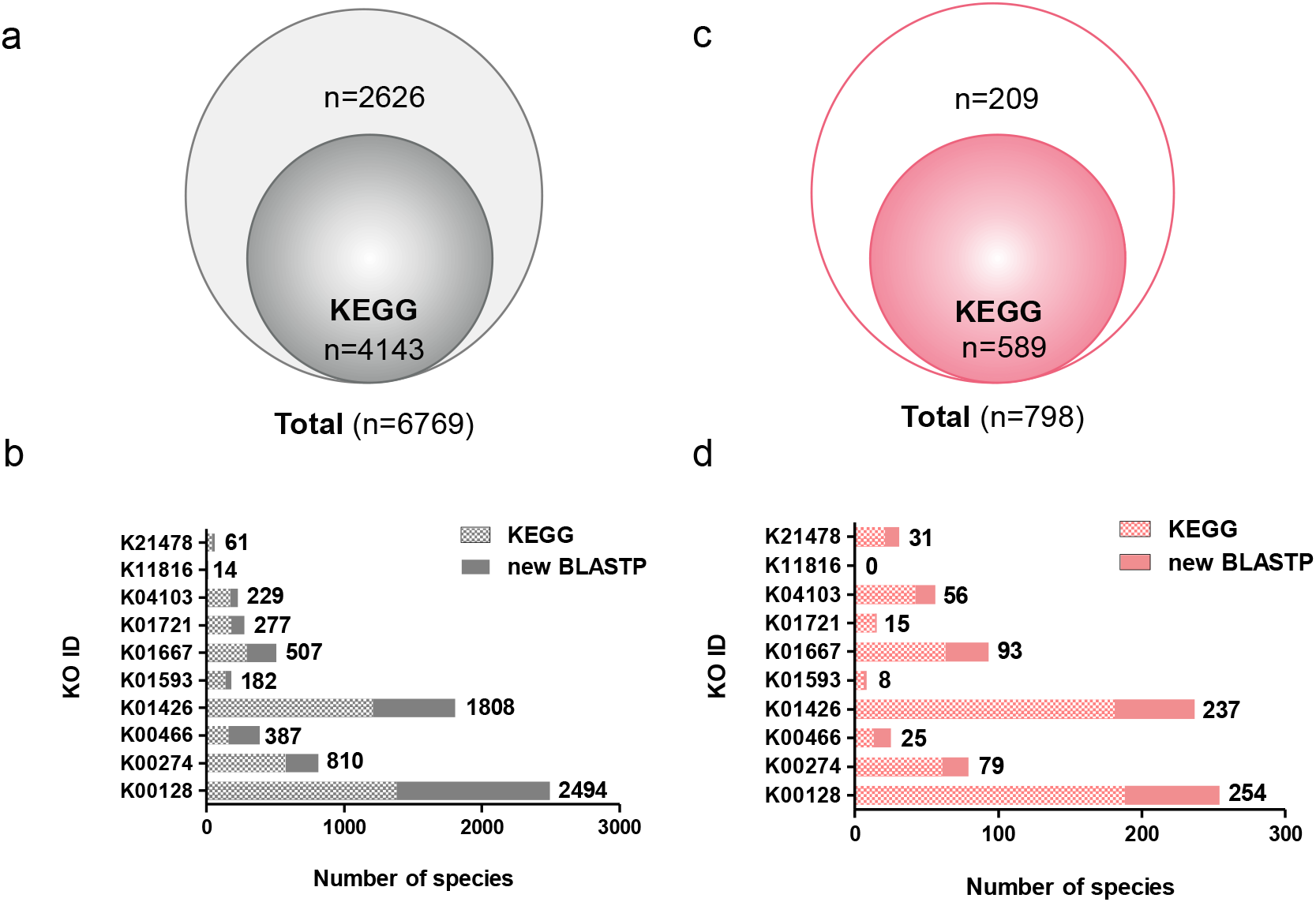
Prediction of new species involved in tryptophan metabolism using BLASTP. Venn diagrams representing the number of prokaryotic species containing genes involved in the production of indoles, reported in KEGG or newly predicted by BLASTP, before (a) and after (c) filtering by the gut microbiota. Bar plots showing the number of prokaryotic species involved in each reaction of indole production, reported in KEGG or newly predicted by BLASTP, before (b) or after (d) filtering for gut microbiota.

We then applied a filtering strategy to select prokaryotic species present in the human gut (**Figure 2c-d**). For this purpose, we constructed a database including gut prokaryote species described in HMP and cultivation studies (Rajilic-Stojanovic and de Vos 2014, Browne, Forster et al. 2016, Lagier, Khelaifia et al. 2016), comprising 1,859 organisms from 1,604 species. In the gut-filtered dataset, BLASTP searches predicted a total of 209 novel prokaryotic species (**Figure 2c**). While some reactions are performed by a reduced number of species (e.g. “K01593”, *n* = 8 and “K01721”, *n* = 15; corresponding respectively to 0.8 % and 1.7 % of the gut prokaryotic dataset), others (e.g. “K00128”) are predicted to be performed by 16.9 % of the gut prokaryotes (*n* = 254) (**Figure 2d and Table 1**). No gut prokaryotic species were predicted to be able to catalyse the conversion of indolepyruvate into indole-3-acetate.

**Table 1.** Microbial genes and species involved in indole production from tryptophan.

| Reaction ID | KO ID | KO Name | Prokaryotic species |  | Gut prokaryotic species |  |
| --- | --- | --- | --- | --- | --- | --- |
|  |  |  | KEGG | + BLASTP | KEGG | + BLASTP |
| **R02678 / R04903** | **K00128** | **ALDH** | **1380** | **1114** | **188** | **66** |
| R02173 / R02908 | K00274 | MAO | 574 | 236 | 61 | 18 |
| **R00679** | **K00466** | **iaaM** | **159** | **228** | **13** | **12** |
| R03096 | K01426 | amiE | 1207 | 601 | 181 | 56 |
| **R00685** | **K01593** | **TDC** | **136** | **46** | **6** | **2** |
| R00673 | K01667 | tnaA | 291 | 216 | 63 | 30 |
| **R04020** | **K01721** | **nthA** | **181** | **96** | **14** | **1** |
| R01974 | K04103 | ipdC | 171 | 58 | 42 | 14 |
| **R10181** | **K11816** | **YUCCA** | **7** | **7** | **0** | **0** |
| R03096 | K21478 | icaB | 37 | 24 | 21 | 10 |

Microbial reactions, genes and number of species involved in the production of indolic compounds from tryptophan, reported in the KEGG orthology database and newly identified by BLASTP searches (+BLASTP).

### Prokaryotic communities involved in the production of indole and indole-3-acetate

Having provided an overview of the prokaryotic metabolism of indoles, we then focused on specific indolic compounds reported to be relevant for human health: indole and indole-3-acetate.

Indole is synthesised in the gut upon the action of a prokaryotic lyase, tryptophanase (“K01667”), followed by conversion by hepatic enzymes into indoxyl and 3-indoxylsulphate (**Figure 3a**). In total, 93 prokaryotic species are involved in this metabolic conversion, amongst which 30 were newly predicted by BLASTP (**Figures 2d** and **Table 1**). Data visualization of these species, in the form of a phylum-clustered network, reveals a marked relative contribution of the phyla *Proteobacteria* and *Bacteroidetes* (**Figure 3a** and **Supplementary table 1**).

**Figure 3.**
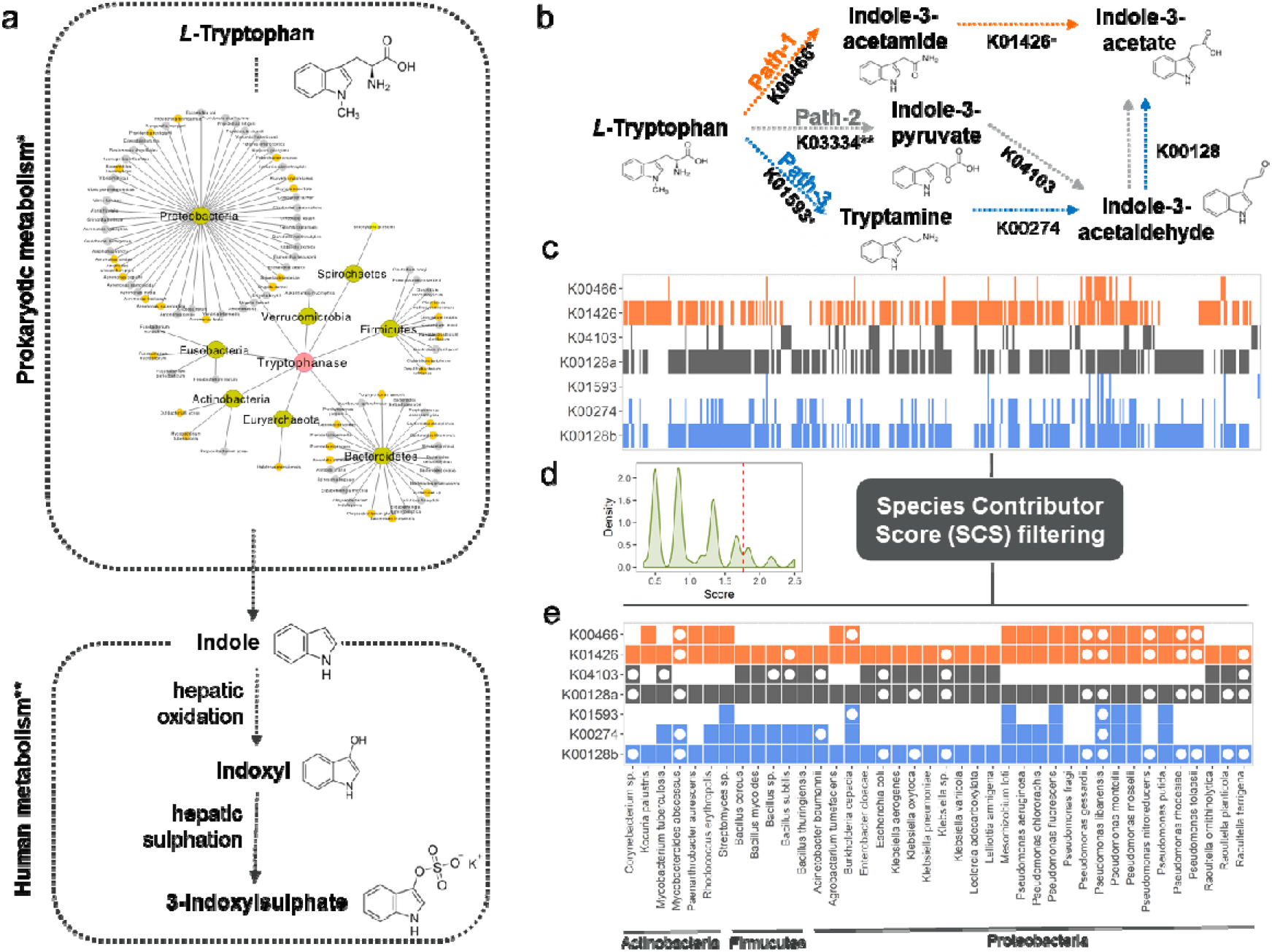
Examples of the contribution of the gut microbiota to the production of indoles. (a) Network representing the gut microbial species involved in the production of indole via tryptophanase, where species newly predicted by BLASTP are colored in yellow. (d) Schematic representation of the 3 different metabolic paths leading to indole-3-acetate. (c) Heatmap showing the contribution of individual species to each of the reactions involved in the production of indole-3-acetate. (d) Histogram showing the distribution of SCS values. (e) Filtered heatmap including species with SCS > 1.66. ‘Exclusively bacterial reactions, “exclusively human reactions.

The production of indole-3-acetate exhibits considerable metabolic complexity, occurring through three metabolic paths: via indole-3-acetamide (path-1, exclusively bacterial), via indole-3-pyruvate (path-2) or via tryptamine (both microbial and mammalian co-metabolism) (**Figure 3b**). An overview of the contribution of prokaryotic species to the individual reactions of the three paths is shown in **Figure 3c**. For each path, the first step is consistently predicted to be the one performed by a lower number of prokaryotic species, suggesting the critical role of a subset of species on the initiation of these metabolic paths.

Prokaryotic species implicated in several steps of the same path, or even in different paths, are likely to be particularly relevant in the synthesis of indole-3-acetate. In order to highlight the most relevant species, we used a scoring system that reflects the overall contribution of each species to these three metabolic paths (**Figure 3d**). We then filtered the original heatmap (**Figure 3c**) to include species with a SCS > 1.66, which corresponds to the ability to perform at least four of the seven reactions (**Figure 3e**). Overall, the species with highest SCS belong to the phylum Proteobacteria. The genus *Pseudomonas* exhibited particularly high SCSs, with five species *(P. putida, P. monteilii, P. mosselii, P. fluorescens, P. libanensis)* predicted to be able to perform six out of the seven metabolic reactions involved in the production of indole-3-acetate (**Figure 3e**). While most of these species were reported in KEGG, BLASTP allowed the identification of several novel species with high SCSs, such as *Pseudomonas* spp. *(P. libanensis, P. nitroreducens, P. rhodesiae, P. tolaasii, P. gessardii), Mycobacteroides abscessus*, and *Raoultella terrigena*.

## DISCUSSION

Unlike previous studies, we used a systematic computational approach to predict the contribution of the gut microbiota to indole metabolism. The production of most indoles (e.g. indole and indole-3-acetate) is dependent, at least partially, on microbial metabolism. In contrast, indole-3-pyruvate may be produced by human metabolism only, upon the action of the L-amino-acid oxidase; a secreted enzyme encoded by the gene IL4I1 and expressed in the gastrointestinal tract, particularly in the colon and rectum (Ponten, Jirstrom et al. 2008). The enzyme is released into the intestinal lumen (Ponten, Jirstrom et al. 2008), where it converts tryptophan into indole-3-pyruvate, which can then be used as a substrate for gut prokaryotic metabolism. Therefore, although indole-3-pyruvate is not a gut microbial metabolite *per se*, it can be used as a substrate for the production of microbial metabolites, exemplifying an interesting case of mammalian–microbial metabolic cooperation.

Our proposed BLASTP-enhanced strategy, followed by gut microbiota filtering, allowed the prediction of 209 novel species involved in the metabolic conversion of tryptophan into indolic compounds. For the production of indole, we predicted 75 novel species that are not reported in KEGG.

Using phylum-level analysis, we predicted the contribution of individual phyla for the production of different indoles. While mostly *Proteobacteria* and *Bacteroidetes* cooperate in the single metabolic step involved in the conversion of tryptophan to indole, for indole-3-acetate, prokaryotic species (belonging to the phyla *Proteobacteria, Actinobacteria* and *Firmicutes)* contribute at a multi-step cooperative level. However, it is important to note that the detection of a protein sequence in a microbe’s genome is only an indicator that it may produce a given metabolite from a substrate; it does not guarantee functionality and further *in vitro* studies are needed to confirm metabolic activity. Validation studies should include qualitative (e.g. Kovac’s reagent or Ehrlich’s reagent (Lombard and Dowell 1983) or quantitative assessment (e.g. Liquid Chromatography-Mass Spectrometry (LC-MS) (Badenoch-Jones, Summons et al. 1982)) of indole in pure cultures of bacteria, after growth in sterile tryptophan-containing medium. Nonetheless, our data-mining strategy represents a novel computational approach to understand the relative contribution, at both species- and phyum-level, of the gut microbiota for single metabolic reactions and cooperative metabolic processes relevant for the production of indoles – and to rank and select the best candidate species for validation studies. Since this approach can be easily extended to other compounds, we propose it as a strategy for those interrogating the bacterial metabolism of gut microbial signalling metabolites relevant for human health, in culture collection and metagenomic studies.

## AUTHOR CONTRIBUTIONS

ALN, ARM and LH conceived the idea of the study. ARM, ALN, RA and LH performed the analysis, with meaningful contributions from AU and JC. JMP contributed to the development and implementation of the scoring system. MED, JKN and LH supervised the work. ALN and ARM drafted and finalised the paper, with input from all authors.

## ACKNOWLEGMENTS

This work was supported by PhD studentships awarded to ALN (Portuguese Foundation for Science and Technology (SFRH/BD/52036/2012) and ARM (Medical Research Doctoral Training Centre (MR/K501281/1), Imperial College (EP/M506345/1) and La Caixa Foundation). JMP holds a Rutherford Fund Fellowship at Health Data Research UK (MR/S004033/1). MED’s lab is funded by EU-FP7 METACARDIS (HEALTH-F4-2012-305312) and the UK Medical Research Council (MR/M501797/1). LH was an MRC Intermediate Research Fellow in Data Science (MR/L01632X/1). This work used the computing resources of the UK MEDical BIOinformatics partnership – aggregation, integration, visualisation and analysis of large, complex data (UK MED-BIO), which was supported by the Medical Research Council (grant number MR/L01632X/1).

## REFERENCES

Altschul, S.F., et al. (1990) Basic local alignment search tool, J Mol Biol, 215, 403–410.

Attwood, G., et al. (2006) Production of indolic compounds by rumen bacteria isolated from grazing ruminants, J Appl Microbiol, 100, 1261–1271.

Badenoch-Jones, J., et al. (1982) Mass spectrometric quantification of indole-3-acetic Acid in Rhizobium culture supernatants: relation to root hair curling and nodule initiation, Appl Environ Microbiol, 44, 275–280.

Bales, J. R., et al (1984). Use of high-resolution proton nuclear magnetic resonance spectroscopy for rapid multi-component analysis of urine. Clin Chem 30(3): 426–432.

Barreto, F.C., et al. (2009) Serum Indoxyl Sulfate Is Associated with Vascular Disease and Mortality in Chronic Kidney Disease Patients, Clin J Am Soc Nephro, 4, 1551–1558.

Bouatra, S., et al. (2013) The human urine metabolome, PLoS One, 8, e73076.

Browne, H.P., et al. (2016) Culturing of ‘unculturable’ human microbiota reveals novel taxa and extensive sporulation, Nature, 533, 543–546.

Chamberlain, S., et al. (2018) taxize: Taxonomic information from around the web. R package version 0.9.3.

Chilloux, J., et al. (2016) The microbial-mammalian metabolic axis: a critical symbiotic relationship, Curr Opin Clin Nutr Metab Care, 19, 250–256.

Danaceau, J.P., et al. (2003) A liquid chromatographic-tandem mass spectrometric method for the analysis of serotonin and related indoles in human whole blood, J Anal Toxicol, 27, 440–444.

Du, Y.-L., Alkhalaf, L.M. and Ryan, K.S. (2015) In vitro reconstitution of indolmycin biosynthesis reveals the molecular basis of oxazolinone assembly, Proc Natl Acad Sci U S A, 112, 2717–2722.

Duranton, F., et al. (2012) Normal and pathologic concentrations of uremic toxins, J Am Soc Nephrol, 23, 1258–1270.

Faure, V., et al. (2006) Elevation of circulating endothelial microparticles in patients with chronic renal failure, J Thromb Haemost, 4, 566–573.

Holmes, E., et al (2012). Therapeutic modulation of microbiota-host metabolic interactions. Science Transl Med 4(137): 137rv136.

Hornemann, U., et al. (1971) The biosynthesis of indolmycin, J Am Chem Soc, 93, 3028–3035.

Jean, M. and DeMoss, R.D. (1968) Indolelactate dehydrogenase from Clostridium sporogenes, Can J Microbiol, 14, 429–435.

Kanehisa, M. and Goto, S. (2000) KEGG: kyoto encyclopedia of genes and genomes, Nucleic Acids Res, 28, 27–30.

Lagier, J.-C., et al. (2016) Culture of previously uncultured members of the human gut microbiota by culturomics, Nat Microbiol, 1, 16203.

Lombard, G.L. and Dowell, V.R., Jr. (1983) Comparison of three reagents for detecting indole production by anaerobic bacteria in microtest systems, J Clin Microbiol, 18, 609–613.

Mohammed, N., Onodera, R. and Or-Rashid, M.M. (2003) Degradation of tryptophan and related indolic compounds by ruminal bacteria, protozoa and their mixture in vitro, Amino Acids, 24, 73–80.

Nicholson, J. K., et al. Gut microorganisms, mammalian metabolism and personalized health care. Nat Rev Microbiol 3(5): 431–438.

Ponten, F., Jirstrom, K. and Uhlen, M. (2008) The Human Protein Atlas - a tool for pathology., J Pathol, 387–393.

Rajilic-Stojanovic, M. and de Vos, W.M. (2014) The first 1000 cultured species of the human gastrointestinal microbiota, FEMS Microbiol Rev, 38, 996–1047.

Roberts, J. and Rosenfeld, H.J. (1977) Isolation, crystallization, and properties of indolyl-3-alkane alpha-hydroxylase. A novel tryptophan-metabolizing enzyme, J Biol Chem, 252, 2640–2647.

Rodriguez-Martinez, A., et al. (2017) MetaboSignal: a network-based approach for topological analysis of metabotype regulation via metabolic and signaling pathways, Bioinformatics, 33, 773–775.

Russell, W.R., et al. (2013) Major phenylpropanoid-derived metabolites in the human gut can arise from microbial fermentation of protein, Mol Nutr Food Res, 57, 523–535.

Shannon, P., et al. (2003) Cytoscape: a software environment for integrated models of biomolecular interaction networks, Genome Res, 13, 2498–2504.

Speedie, M.K., Hornemann, U. and Floss, H.G. (1975) Isolation and characterization of tryptophan transaminase and indolepyruvate C-methyltransferase. Enzymes involved in indolmycin biosynthesis in Streptomyces griseus, J Biol Chem, 250, 7819–7825.

Sun, J.C., et al. (2013) Evaluating effects of penicillin treatment on the metabolome of rats, J Chromatogr B, 932, 134–143.

Takai, K., et al. (1977) Crystalline hemoprotein from Pseudomonas that catalyzes oxidation of side chain of tryptophan and other indole derivatives, J Biol Chem, 252, 2648–2656.

Turnbaugh, P.J., et al. (2007) The human microbiome project, Nature, 449, 804–810.

Venkatesh, M., et al. (2014) Symbiotic bacterial metabolites regulate gastrointestinal barrier function via the xenobiotic sensor PXR and Toll-like receptor 4, Immunity, 41, 296–310.

Wesoly, R. and Weiler, U. (2012) Nutritional Influences on Skatole Formation and Skatole Metabolism in the Pig, Animals (Basel), 2, 221–242.

Whitehead, T.R., et al. (2008) Catabolic pathway for the production of skatole and indoleacetic acid by the acetogen Clostridium drakei, Clostridium scatologenes, and swine manure, Appl Environ Microbiol, 74, 1950–1953.

Wikoff, W.R., et al. (2009) Metabolomics analysis reveals large effects of gut microflora on mammalian blood metabolites, Proc Natl Acad Sci U S A, 106, 3698–3703.

Yang, C.-Y. and Tarng, D.-C. (2018) Diet, gut microbiome and indoxyl sulphate in chronic kidney disease patients, Nephrology (Carlton), 23 Suppl 4, 16–20.

Yisireyili, M., et al. (2013) Indoxyl sulfate promotes cardiac fibrosis with enhanced oxidative stress in hypertensive rats, Life Sci, 92, 1180–1185.

